# EBV LMP1-activated mTORC1 and mTORC2 Coordinately Promote Nasopharyngeal Cancer Stem Cell Formation

**DOI:** 10.1101/2021.11.10.468172

**Authors:** Nannan Zhu, Qian Wang, Zhidong Wu, Yan Wang, Mu-Sheng Zeng, Yan Yuan

**Author notes:** Corresponding authors. Yan Yuan, Department of Basic and Translational Sciences, University of Pennsylvania School of Dental Medicine, Philadelphia, PA 19104.; Mu-Sheng Zeng, State Key Laboratory of Oncology in South China, Sun Yat-sen University Cancer Center, Guangzhou 510060, China.

## Abstract

Epstein-Barr Virus (EBV) is associated with several malignant diseases, including Burkitt’s lymphoma, nasopharyngeal carcinoma (NPC), certain types of lymphomas, and a portion of gastric cancers. Virus-encoded oncoprotein LMP1 induces the epithelial-to-mesenchymal transition (EMT), leading to cancer stem cell formation. In the current study, we investigated how LMP1 contributes to cancer stem cell development in NPC. We found that LMP1 plays an essential role in acquiring CSC characteristics, including tumor initiation, metastasis, and therapeutic resistance by activating the PI3K/mTOR/Akt signaling pathway. We dissected the functions of distinct signaling (mTORC1 and mTORC2) in the acquisition of different CSC characteristics. Side population (SP) formation, which represents the chemotherapy resistance feature of CSC, requires mTORC1 signaling. Tumor initiation capability is mainly attributed to mTORC2, which confers on NPC the capabilities of proliferation and survival by activating mTORC2 downstream genes c-Myc. Both mTORC1 and mTORC2 enhance cell migration and invasion of NPC cells, suggesting that mTORC1/2 co-regulate metastasis of NPC. The revelation of the roles of the mTOR signaling pathways in distinct tumorigenic features provides a guideline for designing efficient therapies by choosing specific mTOR inhibitors targeting mTORC1, mTORC2, or both to achieve durable remission of NPC in patients.

**Significance:** LMP1 endows NPC to gain cancer stem cell characteristics through activating mTORC1 and mTORC2 pathways. The different mTOR pathways are responsible for distinct tumorigenic features. Rapamycin-insensitive mTORC1 is essential for CSC drug resistance. NPC tumor initiation capacity is mainly attributed to mTORC2 signaling. mTORC1 and mTORC2 co-regulate NPC cell migration and invasion. The revelation of the roles of mTOR signaling in NPC CSC establishment has implications for novel therapeutic strategies to treat relapsed and metastatic NPC and achieve durable remission.

## Introduction

Nasopharyngeal carcinoma (NPC) is one of the most aggressive head and neck malignancy arising from the nasopharynx epithelium. NPC has unique geographic distribution and affects defined populations, mainly in Southern China, Southeast Asia, the Arctic, and Northern Africa (1). In these endemic regions, most NPC cases (>95%) are non-keratinizing carcinoma and invariably associated with Epstein-Barr virus (EBV) infection (2). Although NPC patients’ overall survival has improved in recent years, 21.3% of patients in a study suffered from recurrence, distant metastases and radiotherapy failure (3). Cancer stem cells (CSCs) have been implicated to be involved in cancer relapse and metastasis. Understanding how CSCs are generated and maintained in NPC will lead to novel strategies for NPC treatment.

EBV latent membrane protein1 (LMP1) is an oncoprotein and detected in most of invasive and malignant NPC lesions (4). It has been shown that LMP1 induces the epithelial-to-mesenchymal transition (EMT) and increase metastasis of NPC (5). We reported that LMP1 plays a crucial role in promoting EMT in NPC to generate various subpopulations of CSCs arrayed along the epithelial (E) to mesenchymal (M) spectrum. Furthermore, we demonstrated that the hybrid E/M state exhibits the highest tumor initiating capacity, while the xM state contributes to vasculogenic mimicry, a hallmark of metastatic cancers (6). However, the mechanism underlying LMP1 regulating CSC development and maintenance remains unknown. Transcriptomic analysis revealed that the PI3K/mTOR/Akt signaling is the most significantly affected pathway in EBV-infected NPC cells compared to EBV negative NPC cells (7). This finding, together with the previous report that LMP1 activates the mTOR/AKT signaling pathway (8, 9), prompted us to investigate if the mTOR signaling pathway plays a role in EMT and CSC development in NPC.

The mammalian target of rapamycin (mTOR) is the principal regulator of growth controlling cell proliferation, survival, anabolic and catabolic processes in response to nutrients and environment (10). mTOR exists in two structurally and functionally distinct protein complexes, namely mTOR complex C1 (mTORC1) that is highly sensitive to rapamycin, and mTOR complex 2 (mTORC2) that is resistant to short-term treatment of rapamycin (11-13). Both mTORC1 and mTORC2 share several common subunits: the mTOR kinase, mLST8, Deptor, and Tti1/2. Additionally, each complex has distinct subunits: Raptor and PRAS40 are subunits specific to mTORC1 (12), while Rictor and mSin1 are unique to mTORC2 (13, 14). mTORC1 is stimulated by PI3K/AKT and Ras-MAPK cascades and, once activated, phosphorylates EIF-4B binding protein 1(4E-BP1) and S6 kinase1 (S6K1) to trigger cellular proliferation (11, 12). Although the function and regulation of mTORC2 are not as well-defined as mTORC1, sufficient evidence suggests that mTORC2 plays fundamental roles in regulating cell metabolism (15, 16). mTORC2 phosphorylates several AGK kinase family members, including AKT (Ser473), PKCα(Ser638, and Ser657)and SGK (17). AKT integrates signals from mTORC2 (Ser473) and from PDK1 (Ser318) to promote cell growth and survival and is among the most commonly hyper-activated proteins in cancers (18). As mTOR signaling is a chief mechanism for controlling cell proliferation, survival and metabolism, it is not surprising that mTOR signaling is one of the most frequently activated pathways in cancer (19). Accumulating evidence demonstrated the involvement of mTOR signaling in cancer initiation, metastasis and resistance to cancer therapies, suggesting its role in EMT (20, 21).

In this study, we investigated the contribution of the mTOR pathway to the generation and maintenance of CSCs in NPC. shRNA-mediated gene silencing approach and mTOR complex-specific pharmacological inhibitors were used to dissect the function of mTOR or its complexes in each step of the EMT process that leads to the development of nasopharyngeal cancer stem cells. Our study indicates that the mTOR pathways are essential for NPC to maintain CSCs and tumorigenicity, and mTORC1 and mTORC2 play distinct roles in different steps of CSC development.

## Results

### LMP1 activates the mTOR signaling pathway

Our previous study showed that LMP1 could induce epithelial-to-mesenchymal-transition (EMT) in NPC, resulting in the generation of cancer stem cells at various states along the epithelial– mesenchymal spectrum and the acquisition of tumorigenic and metastatic capabilities (6). The LMP-induced EMT process is accompanied by activation of mTORC1 and mTORC2 pathways revealed by the up-regulation of mTORC1 and mTORC2 in LMP1-induced epithelial/mesenchymal (E/M) hybrid and highly mesenchymal (xM) subpopulation (Fig. 1A). This finding, together with the previous report that LMP1 regulates the mTOR signaling pathway in NPC (8), prompted us to investigate if LMP1 induces the EMT program and cancer cell stemness by activating the mTOR pathway. To this end, we first examined the effect of LMP1 expression on the activation status of mTOR components in CNE-2 S26 (S26) and CNE-2 S18 (S18) cells, two clones derived from the same NPC cell line CNE-2 but with different epithelial-mesenchymal phenotypic states. S18 displays mesenchymal-like (M-like) phenotypes and possesses great metastasis abilities, while S26 shows epithelial-like (E-like) phenotypes and produces low invasion and metastasis (22). In both S18 and S26 cells, transient expression of LMP1 resulted in elevated phosphorylation of mTOR (S2448) and p70S6K (T389) in the mTORC1 pathway in a dose-dependent manner. Furthermore, LMP1 also promoted the phosphorylation of substrates of mTORC2, including PKCα (S638) and Akt (S473) (Fig. 1B). In addition, stably expressing LMP1 in S26 and S18 increased phosphorylation of key effectors of mTORC1 and mTORC2, including Akt (S473), c-Raf (S259), and mTOR (S2448) (Fig. 1C). Overall, LMP1 induces the activation of mTORC1 and mTORC2 in NPC cells, suggesting an essential role of mTOR signaling in the regulation of cancer stem cell formation.

**Figure 1.**
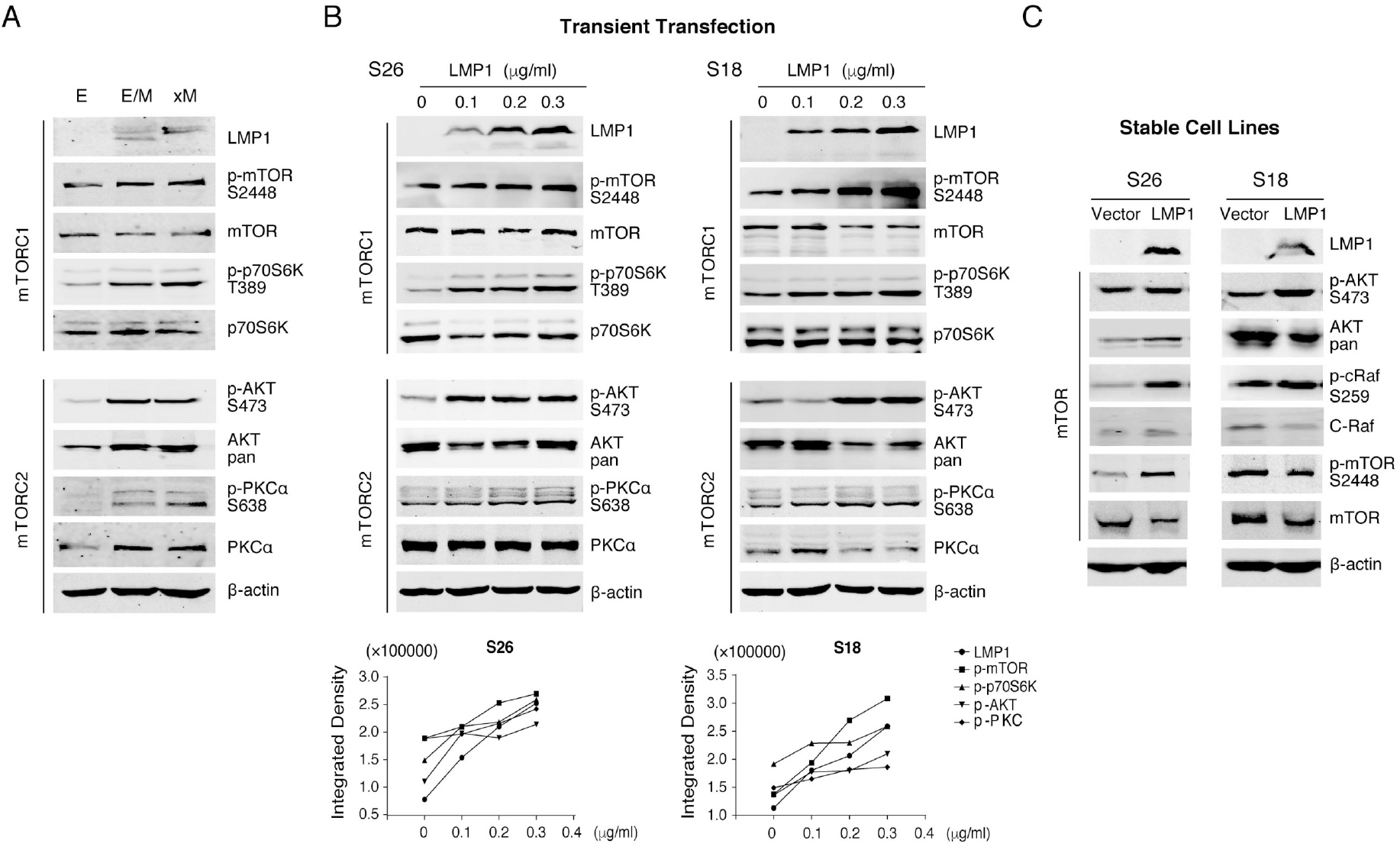
Effect of LMP1 expression on mTOR signaling. (**A)** E/M and xM subpopulations sorted from S26-LMP1/2A stable cell line, and E subpopulation sorted from S26 cells were analyzed by Western blotting for the expression and phosphorylation of mTOR components and substrates. (**B)** CNE-2 S26 and S18 cells were transfected with various doses of pcDNA3.1-LMP1. Forty-eight hours post-transfection, cell lysates were analyzed by Western blotting for the expression and phosphorylation of mTOR components and substrates. The expression or phosphorylation of proteins was quantitated by densitometry and plotted. (**C)** S26 and S18 cells that stably express LMP1 were examined by Western blotting for activation of mTOR sigaling. β-actin serves as the loading control.

### LMP1-mediated activation of mTORC1 is responsible for cancer stemness properties of NPC cells

Cancer stem cells can be identified or isolated based on specific surface and enzyme markers and characterized by self-renewal ability, differentiation into non-CSC progeny tumor cells, high tumorigenicity, and resistance to chemotherapy. To define the roles of the mTOR pathway in NPC cells for the acquisition of stemness and CSC characteristics, we examined the contributions of mTORC1 and mTORC2 complexes to each of the CSC properties. First, we determined the effects of the mTOR pathways on cancer cell stemness by following the changes of mesenchymal marker Vimentin and cancer stem cell marker aldehyde dehydrogenase (ALDH) in response to the mTORC1 and mTORC2 signalings. We employed a loss-of-function strategy to determine if mTOR pathways are responsible for expressing the stem cell markers. A short-hairpin RNA (shRNA) against mTOR was introduced into S18 cells to silence mTOR gene expression. Also, the functions of mTORC1 and mTORC2 were respectively blocked by using shRNAs specific to mTOR complex-specific components Raptor (mTORC1) and Rictor (mTORC2). The efficiency of each shRNA-mediated gene silencing, and the functional consequence such as the phosphorylation of p70S6K at T389 (a downstream effector of mTORC1) and AKT at S473 (a core kinase of mTORC2 signaling), were analyzed by Western blot. As shown in Fig. 2A, these shRNAs effectively inhibited the expression of the targeting mTOR components. The consequence of mTORC1 and mTORC2 knockdown on Vimentin expression was assessed by Western analysis and IFA. Results showed that loss of mTORC1 function significantly reduced the mesenchymal marker vimentin (Fig. 2B and C).

**Figure 2.**
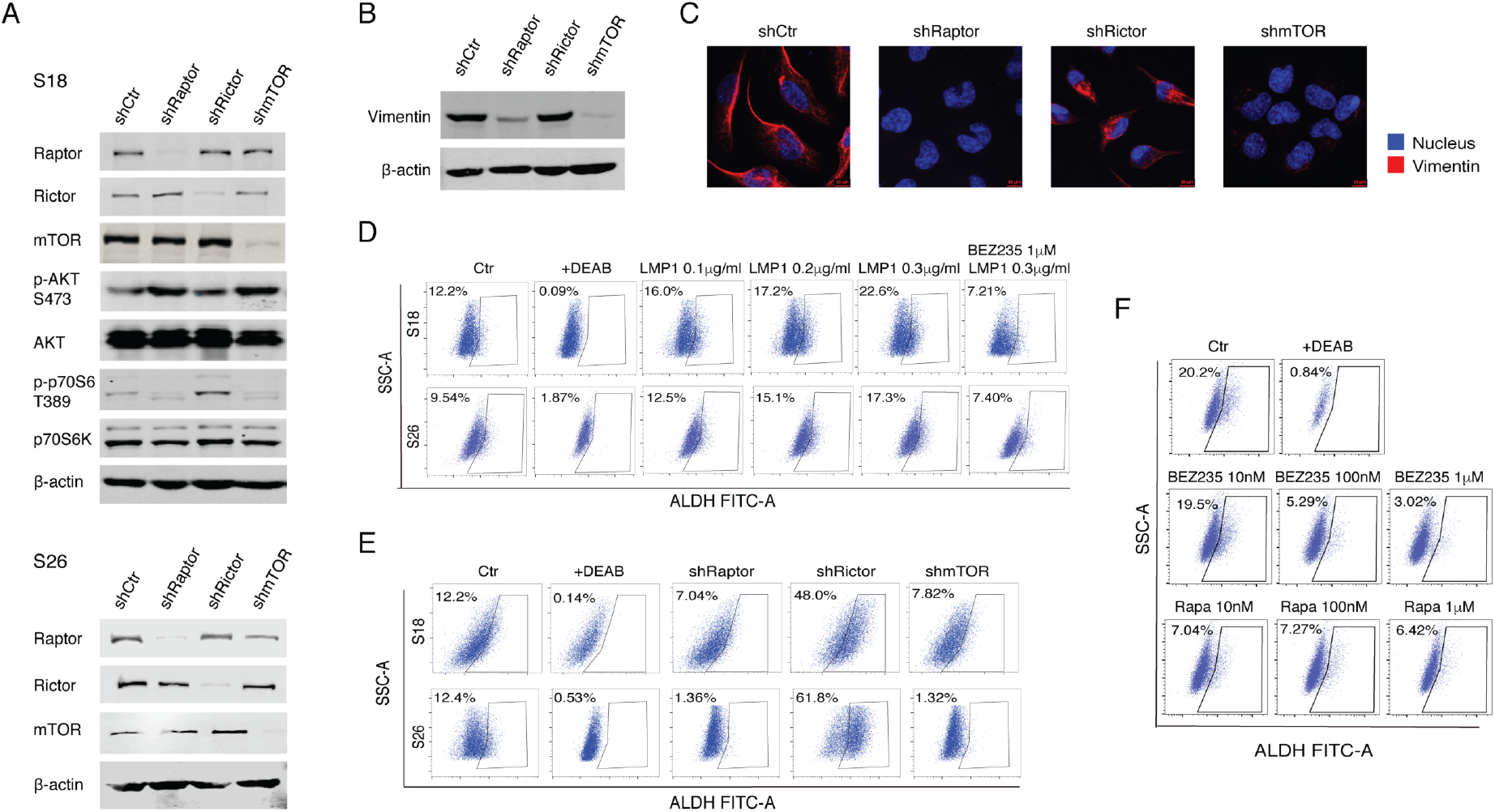
LMP1-mediated activation of mTORC1 confers on NPC cells cancer stemness properties. (**A)** CNE-2 S18 cells were transduced with shRNA lentiviruses against mTOR, Raptor, and Rictor genes. The knockdown efficiencies and the functional consequences (phosphorylation of AKT at S473 and p70 S6K at T389) were analyzed by Western blots. (**B and C)** The effect of silencing mTOR signaling in S18 cells on Vementin expression was analyzed by Western **(B)** and immunofluorescent assays (630X) **(C). (D)** CNE-2 S26 and S18 cells were transfected with pcDNA3.1-LMP1 or combination of 0.3μg/ml pcDNA3.1-LMP1 and 1μM BEZ235. Forty-eight hours post-transfection, cells were subjected to flow cytometry (Aldefluor) for ALDH^br^ cells. ALDH^br^ gating is established using the cells treated with ALDH inhibitor DEAB. **(E)** S18 and S26 cells were transduced with shRaptor, shRictor and shmTOR lentiviruses, selected with puromycin for 7 days, and then analyzed by Aldefluor assay. **(F)** S18 cells were treated with mTORC1/2 dual inhibitor BEZ235 and mTORC1 inhibitor rapamycin for 24 hours. The effects of these treatments were determined by monitoring ALDH^br^ populations using Aldefluor assay.

ALDH is an enzyme for the oxidation of intracellular aldehydes and essential for the maintenance and differentiation of stem/progenitor cells in normal development. ALDH is a hallmark of cancer stem cells and a prognostic factor of poor clinical outcomes (23, 24). Cells expressing a high level of ALDH become brightly fluorescent (ALDH^br^), which allows us to determine the percentage of ALDH^br^ cells in S18 and S26 cell populations and the effect of LMP1 expression on ALDH expression using flow cytometry. S18 and S26 consisted of 12.2% and 9.54% ALDH^br^ cells, respectively, which were significantly reduced when the cells were treated with a specific inhibitor of ALDH enzyme, Diethylaminobenzaldehyde (DEAB). LMP1 expression increased the ALDH^br^ population in a dose-dependent manner in S18 (from 12.2% to 22.6%) and S26 cells (from 9.54% to 17.3%), but the LMP1-mediated increases were diminished when the cells were treated with PI3K-mTOR dual inhibitor BEZ235 (Fig. 2D). The involvement of mTOR signaling in the maintenance of ALDH^br^ cells was dissected using shRNA-mediated knockdown of mTOR, Raptor, and Rictor. Results showed that silencing Raptor and mTOR decreased ALDH^br^ cell levels in S18 and S26 cells (Fig. 2E), suggesting mTORC1 is responsible for the acquisition of ALDH^br^ phenotype in NPC. To our surprise, the knockdown of Rictor increased the population of ALDH^br^ cells by nearly 4–5 fold in both S18 and S26 cells (Fig. 2E). A possible explanation is that block of mTORC2 may lead to the feedback of increased mTORC1 as revealed by dramatically elevated phosphorylation of p70S6K, a key effector of mTORC1 (Fig. 2A). Taken together, mTORC1 is essential for maintaining ALDH^br^ CSCs in NPC. This notion was also confirmed by treating S18 cells with mTORC1 inhibitor Rapamycin and PI3K-mTOR dual inhibitor BEZ235, which significantly decreased the ALDH^br^ cell population in S18 (Fig. 2F).

### Rapamycin-insensitive mTORC1 is essential for CSC drug resistance

One crucial characteristic of CSC is the resistance to chemotherapy, represented by the capacity of cells to extrude dyes such as Hoechst33342 due to the high expression of ATP-dependent efflux pumps (25). The cells that extrude the dye and maintain a low fluorescent signal are referred to as side population (SP) cells. SP cells have been identified in NPC (26). S26 clone of CNE-2 was found to possess a low percentage of SP cells (3.56%), while S18 clone has a high population of SP cells (38.7%) (Fig. 3A). Ectopic expression of LMP1 increased the SP cell population in S26 and S18 cells, but such SP increase diminished when the cells were treated with mTOR dual inhibitor BEZ235 (Fig. 3B). These data suggest that LMP1-activated mTOR signaling is crucial for the drug resistance property of CSCs in NPC. To further analyze the role of mTOR and its complexes in developing a drug-resistant NPC population, we knocked down the expression of mTOR, Raptor, and Rictor, respectively (Fig. 3C) and analyzed their effects on the side population. Notably, silencing the expression of either Raptor or mTOR significantly reduced the side population, while knockdown of Rictor exhibited little change in SP (Fig. 3D), suggesting that mTORC1 played an essential role in developing drug resistance of NPC.

**Figure 3.**
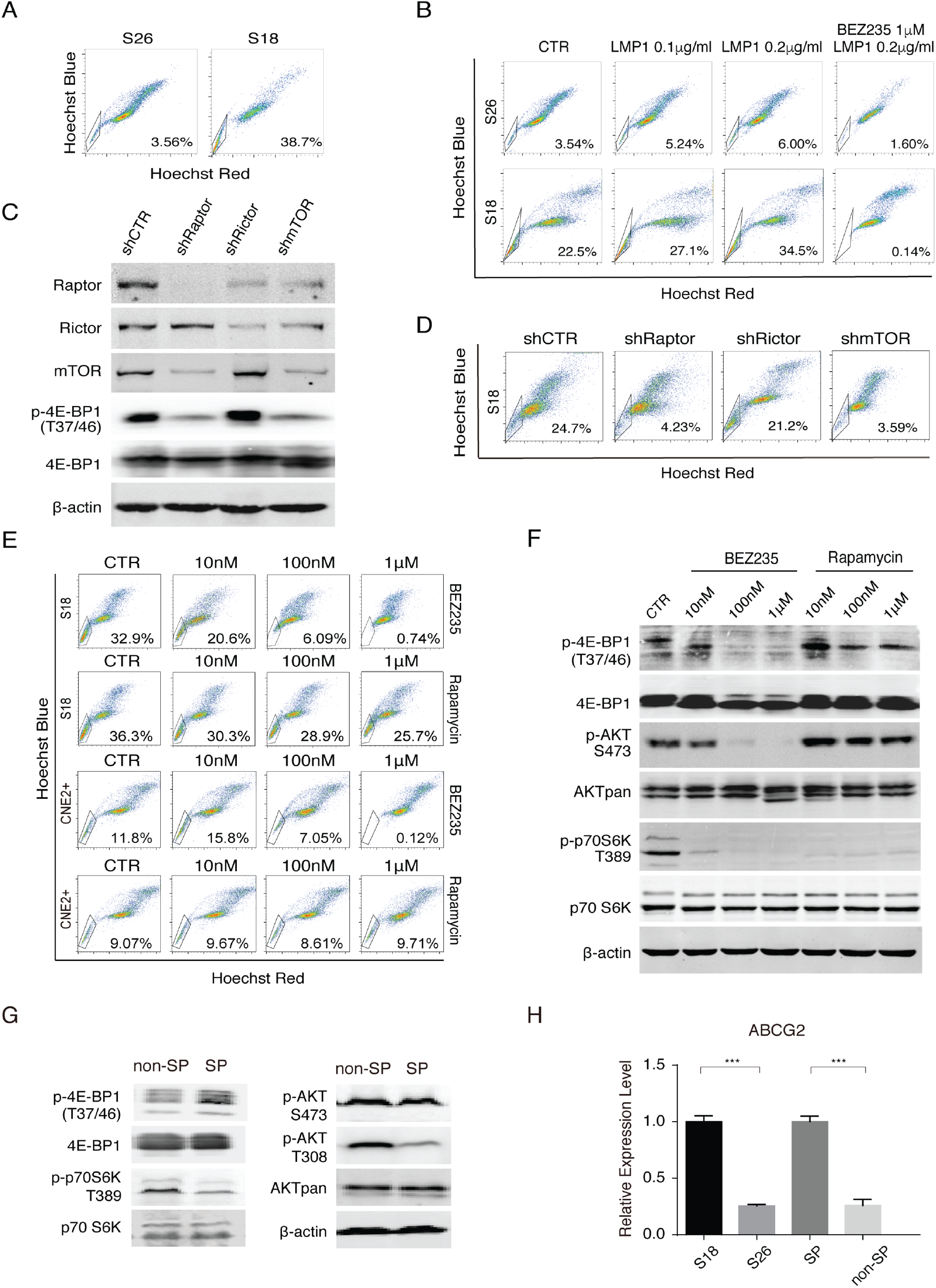
Rapamycin-insensitive mTORC1 is essential for NPC drug resistance. (**A)** Flow cytometry analysis of side population (SP) in CNE-2 S26 cells and S18 cells stained with Hoechst 33342. (**B)** S26 and S18 cells were transfected with pcDNA3.1-LMP1 or treated with combination 1μM BEZ235 and 0.2μg/ml pcDNA3.1-LMP1. 48 hours post-transfection, cells were subjected to flow cytometry analysis for SP cells. (**C)** S18 cells were transduced by specific shRNA lentiviruses to silence the expression of Raptor (shRaptor), Rictor (shRictor) and mTOR (shmTOR), respectively. The knockdown efficiency of each shRNA was verified by Western blot, as well as the functional consequences [phosphorylation of 4E-BP1(T37/46) and 4E-BP1]. (**D)** Effects of shRNA-mediated knockdown of mTOR components were analyzed for SP by flow cytometry. (**E)** S18 cells and EBV-positive CNE2 cells (CNE2+) were treated mTORC1/2 dual inhibitor BEZ235 and mTORC1 inhibitor rapamycin in various doses for 24 hours. The effects of these treatments on SP were analyzed by flow cytometry after staining with Hoechst 33342. **(F)** The effects of the treatment on phosphorylation of 4E-BP1 at T37/46, p70S6K at T389 and AKT at S473 in S18 cells were examined. **(G)** Side population cells (SP) and non-side population cells (non-SP) were sorted from CNE2-S18 cells by FCS assay. The expression and phosphorylation of mTOR components and substrates in SP and non-SP cells were analyzed by Western blots. (H) The expression of ABCG2 mRNA relative to GAPDH in SP, non-SP, S18, and S26 cells was determined by RT-qPCR (Mean +/- SD of three biological replicates).

To verify the conclusion, flow cytometry assays for SP cells were performed in EBV-negative CNE S18 and EBV-positive CNE2 cells (CNE2+) treated with mTORC1 inhibitor Rapamycin and mTORC1/2 dual inhibitor BEZ235. To our surprise, Rapamycin did not effectively decrease side population cells. In contrast, BEZ235 significantly reduced SP cells in a dose-dependent manner (Fig. 3E). It was reported that when the rapamycin-insensitive mTORC1 (RI-mTORC1) complex is activated, rapamycin couldn’t block all downstream effectors of mTORC1. In these cells, although rapamycin decreases the phosphorylation of p70S6K at Thr389, it cannot effectively block the phosphorylation of 4E-BP1 (27). To see if RI-mTORC1 is activated in NPC, S18 cells were treated with rapamycin or BEZ235 and analyzed for phosphorylation of p70S6K and 4E-BP1 by Western blot. Indeed, rapamycin inhibited the phosphorylation of p70S6K, but not 4E-BP1 (Thr37/46) in S18 cells. BEZ235 completely blocked the phosphorylation of both p70S6K and 4E-BP1 (Thr37/46) (Fig. 3F), indicating a RI-mTORC1 activity is responsible for the SP maintenance and drug resistance property of NPC cancer stem cells. We isolated SP and non-SP cells and examined them for RI-mTORC1 activities. Western blots showed that the sorted SP cells indeed exhibited higher levels of phosphorylation of 4E-BP1 than non-SP cells (Fig. 3G).

The molecular basis for side population is known to attribute to the ATP-binding cassette sub-family G member 2 (ABCG2) (28, 29). ABCG2 is a multidrug transporter that protects many tissues against xenobiotic molecules. ABCG2 was also considered a marker for cancer stem cells, contributed to multidrug resistance of tumor cells (30). It has been reported that PI3K/mTOR pathway up-regulates ABCG2 expression to regulate side population phenotype in cancer stem cells (31). We compared SP and non-SP cells for their ABCG2 expression levels and results showed that SP cells expressed higher level of ABCG2 transporter compared to non-SP cells (Fig. 3H). S18 cells, which display a high percentage of side population, also expressed ABCG2 in a higher level compared to S26 that exhibits a low side population. These results indicated that the fundamental difference between SP and non-SP is the expression level of ABCG2, and mTORC1 regulates ABCG2 expression through the RI-mTOR–4E-BP1–ABCG2 axis to enhance the drug-efflux pump activity in NPC.

### mTORC2 is essential for tumor initiation

Another essential characteristic of CSC is tumor initiation capability. Sphere-forming assay is often used to identify CSCs and study their tumor initiation property (32), which allows us to investigate how mTOR pathway participates in regulation of tumor initiation capability. We previously showed that the expression of LMP1 in S26 cells enhanced tumorsphere formation (6). The LMP-mediated tumorsphere formation diminished when S26-LMP1 cells were treated with mTOR dual inhibitor BEZ235, while mTORC1 inhibitor rapamycin exhibited little effect in this phenotype (Fig. 4A). Tumorsphere-forming assays with S18 cells and EBV-positive TW03 cells (TW03+) cells led to the same conclusion (Fig. 4B). To gain insight into how the mTOR pathways contribute to the establishment and maintenance of the tumor initiation capability, mTOR, Raptor and Rictor expression were silenced in S18 and S26 cells by shRNA-mediated knockdown and analyzed using sphere-forming assay. The result showed that silencing of Rictor and mTOR significantly or even completely abolished NPC tumorsphere formation, indicating that mTORC2 is essential for tumor initiation in NPC cells (Fig. 4C and D).

**Figure 4.**
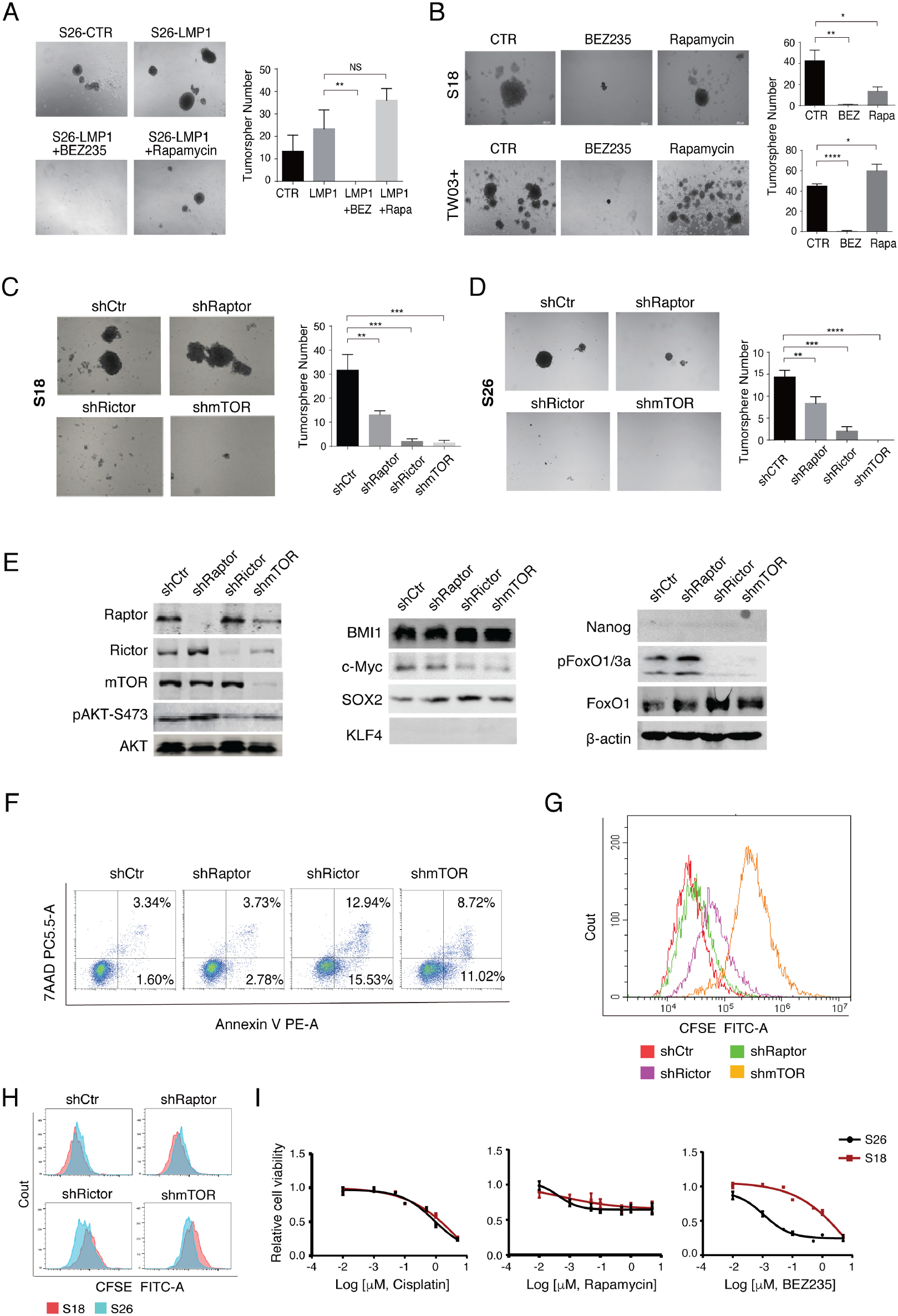
Tumor initiation ability of nasopharyngeal cancer stem cells is mainly attributed to mTORC2. (**A)** CNE2-S26 cells expressing LMP1 were treated with 1μM BEZ235 or 100nM Rapamycin and the effects of mTOR inhibitors on tumor initiation ability were analyzed by tumorsphere-forming assay. Representative images are shown (50X) (mean ±SD, n=3). (**B)** CNE2-S18 cells and EBV-positive TW03 cells (TW03+) were treated with 1μM BEZ235 or 100nM Rapamycin and subjected to tumorsphere-forming assay. Representative images are shown (50X) (mean ±SD, n=3 per group). **(C and D)** S18 cells and S26 cells were transduced with shRNA lentiviruses targeting Raptor (shRaptor), Rictor (shRictor) and mTOR (shmTOR), respectively. The effects of silencing mTOR components on tumor initiation ability were analyzed by tumorsphere-forming assay and representative images are shown (50X). The tumorsphere number of each sample was quantitated (mean ±SD, n=3). (**E)** The knockdown efficiency of each shRNA and functional consequences (the phosphorylation of AKT and FoxO and the expression of pluripotent transcription factors Bmi-1, c-Myc, Sox2, KLF4, and Nanog) in CNE2-S18 cells were examined by Western analysis. (**F)** S18 cells and mTOR component knockdown cells were analyzed for apoptosis by Annexin-V flow cytometry assay. (**G)** S18 and these knockdown cells were subjected to CFSE dye dilution assays for cell proliferation ability. (**H**) Cell proliferation upon the shRNA knockdown of mTOR components were compared between S18 and S26 cells by CFSE dye dilution assays. (**F)** Does-response curves of S18 and S26 cells to the treatment with Cisplatin, Rapamycin and BEZ235 were compared (mean ±SD, n=3).

To elucidate how mTORC2 influenced NPC’s tumor initiation capability, S18 cells and the cells having mTOR, Raptor, and Rictor silenced were analyzed for the expression of several pluripotent transcription factors such as Bmi-1, Sox2, KLF4, Nanog, and c-Myc. These markers have pronounced roles in promoting stemness and driving tumorigenesis and are often used to characterize CSCs in solid tumors (33, 34). Results revealed that the expression of c-Myc was down-regulated in the cells where Rictor and mTOR were silenced, suggesting that mTORC2 may promote tumor initiation capability of NPC cells by regulating oncogene c-Myc (Fig. 4E). c-Myc is highly expressed in cancer stem cells relative to non-stem cells and has functions in self-renewal and differentiation of stem cells (35). The mTORC2/AKT pathway was reported to regulate c-Myc through Forkhead box O (FoxO), a negative regulator of c-Myc (36). When mTORC2/AKT signaling is activated, AKT phosphorylates FOXO on discrete residues, leading to its inactivation and exclusion from the nucleus. Consequently, c-Myc expression is up-regulated (37, 38). To verify if that is the case in NPC, we examined the effect of silencing of mTORC2 on phosphorylation of AKT (Ser473) and FoxO1 (Thr24)/FoxO3a (Thr32) by Western analysis. As shown in Fig. 4E, knockdown of Rictor and mTOR expression led to the down-regulation of Phospho-FoxO1 (Thr24) and FoxO3a (Thr32) as well as AKT (Ser473).

c-Myc activation is a hallmark of cancer initiation and maintenance. When pathologically activated, c-Myc enforces many of the “hallmark” features of cancer, including increased stemness, relentless cellular proliferation, and resistance to apoptosis (39). We examined the effects of the mTORC2–c-Myc axis on NPC cell proliferation and resistance to apoptosis. Using Annexin-V assay, S18 and the Raptor, Rictor and mTOR knockdown cells were analyzed for cell apoptosis. Silencing Rictor and mTOR induced explicitly higher extents of apoptosis of S18 cells (12.94% and 8.72%, respectively) than that of control cells (3.34%) (Fig. 4F). CFSE (carboxyfluorescein diacetate succinimidyl ester) assay was used to measure cell proliferation based on dye-dilution in the daughter cells. Results showed that silencing Rictor or mTOR suppressed S18 cell proliferation compared to silencing Raptor or control vector (Fig. 4G). Additionally, CFSE assay showed that silencing Rictor preferentially inhibited S18 cells (cancer stem-like cells) proliferation rather than S26 cells (Fig. 4H). Furthermore, the use of pharmacological mTOR dual inhibitor BEZ235 also showed that blockade of mTOR signaling preferentially inhibited S18 cell proliferation, while Cisplatin and Rapamycin inhibited S18 and S26 with same efficiencies (Fig. 4I). Taken together, these data suggest that mTORC2 plays an essential role in CSC proliferation and can serve a potential target to selectively inhibit cell proliferation and tumor initiation of nasopharyngeal cancer stem cells.

### mTORC1 and mTORC2 co-regulate migration and invasion

Cancer stem cells are inherently capable of metastasizing. The expression of LMP1 in S26 cells render the low migration/invasion cells dramatically higher abilities of migration and invasion (Fig. 5A). The migration and invasion could be blocked by the treatment with mTORC1/2 dual inhibitor BEZ235 in LMP1-expressing S26 cells and S18 cells, while mTORC1 inhibitor rapamycin showed limited inhibitory effect (Fig. 5A and B). To further explore the role of the mTOR signaling in NPC metastasis, we analyzed S18 cells for the ability of migration and invasion as well as functional consequences of silencing of mTORC1 and mTORC2 functions *in vitro* using wound healing test and Transwell invasion assay. Silencing either Rictor or Raptor could not reduce migration and invasion abilities of S18 cells but knocking down mTOR expression significantly inhibits migration and invasion (Fig. 5C–E). Therefore, these data suggest that mTORC1 and mTORC2 regulate migration and invasion either with some overlapped functions or by coordinating a common signal pathway.

**Figure 5.**
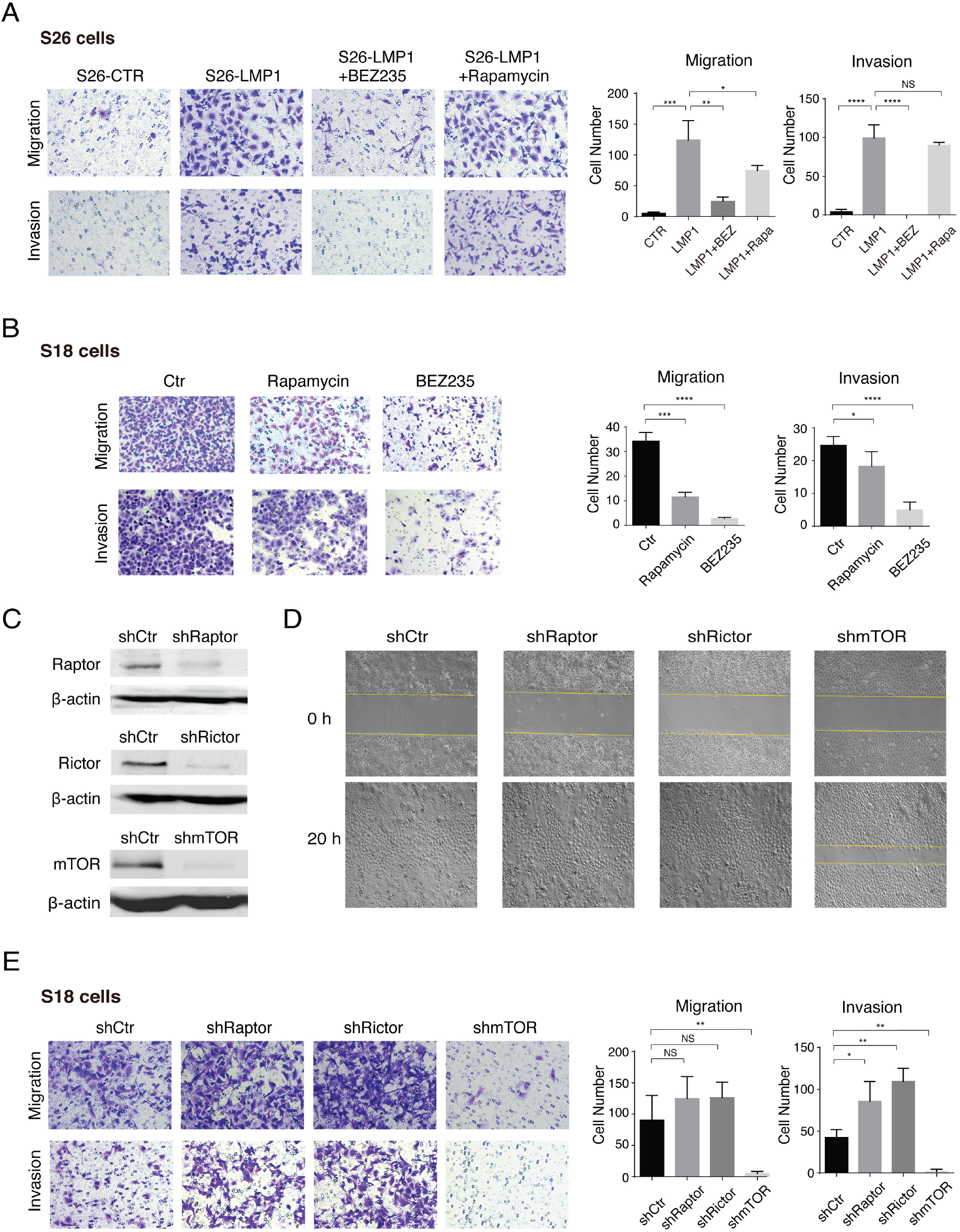
Effects of silencing mTORC1 and mTORC2 on NPC cell migration and invasion capabilities. (**A)** CNE2-S26 LMP1-expressing cells were treated with 1μM BEZ235 or 100nM Rapamycin and effects of the treatments on cell migration and invasion abilities were assayed using Transwell migration and invasion assays. Cells migrated to the lower chamber were fixed, stained with crystal violet, and counted (200X, mean ±SD, n=3). **(B)** CNE2-S18 cells were treated with 1μM BEZ235 or 100nM Rapamycin and analyzed for invasion and migration. The S18 cells that invaded into the lower chamber were fixed, stained and counted (200x, mean ±SD, n=3). (**C and D**) S18 cells and cells expressing shRNAs targeting Raptor (shRaptor), Rictor (shRictor) and mTOR (shmTOR), respectively, were subjected to a wound healing assay for their migration ability. The knockdown efficiencies of shRNAs on each target genes were determined by Western blotting after 7 days of puromycin selection. (**E)** S18 cells and cells expressing shRaptor, shRictor and shmTOR were assayed for their migration and invasion abilities using Transwell migration and invasion assay (200X, mean ±SD, n=3).

### Inhibition of mTOR pathway tampers tumor growth *in vivo*

The essential roles of mTORC1 and mTORC2 pathways in the generation and maintenance of NPC cancer stem cells informs that mTOR can serve as an effective drug target and mTOR inhibitors have potentials to be used to treat advanced NPC. To validate the potential, we tested if blocking the mTOR pathway can inhibit NPC tumor progression *in vivo*. CNE-2 S18 cells and the cells in that mTOR, Raptor, and Rictor were respectively silenced by specific shRNAs were transplanted into BALB/C-nu/nu mice to test their tumorigenic ability *in vivo*. Tumors were stripped after 30 days. We found that silencing either mTORC1 or mTORC2 effectively decreased tumor weight (Fig. 6A). These tumors were analyzed by hematoxylin-eosin (H&E) staining, immunohistochemistry (IHC) assay for Ki67 and TUNEL staining. The expression of Ki67 is strongly associated with tumor cell growth and serves as a proliferation marker. TUNEL staining is used to identify apoptotic cells in tissue based on the labeling of DNA strand breaks. Results showed that silencing mTOR, Raptor and Rictor significantly decreased Ki67 positive cells and increased apoptotic cells (Fig. 6B), suggesting that lack of mTOR function decreased cell proliferation and tumor growth *in vivo*. Additionally, S18-xenograft mice were treated with mTOR dual inhibitor BEZ235 for 21 days. We found that BEZ235 significantly inhibited the growth of formed tumors (Fig. 6C–E) but had little effect on the body weight of the treated mice (Fig. 6F). Taken together, mTORC1 and mTORC2 are required for tumor growth and malignant process, and inhibition of their functions leads to delayed growth.

**Figure 6.**
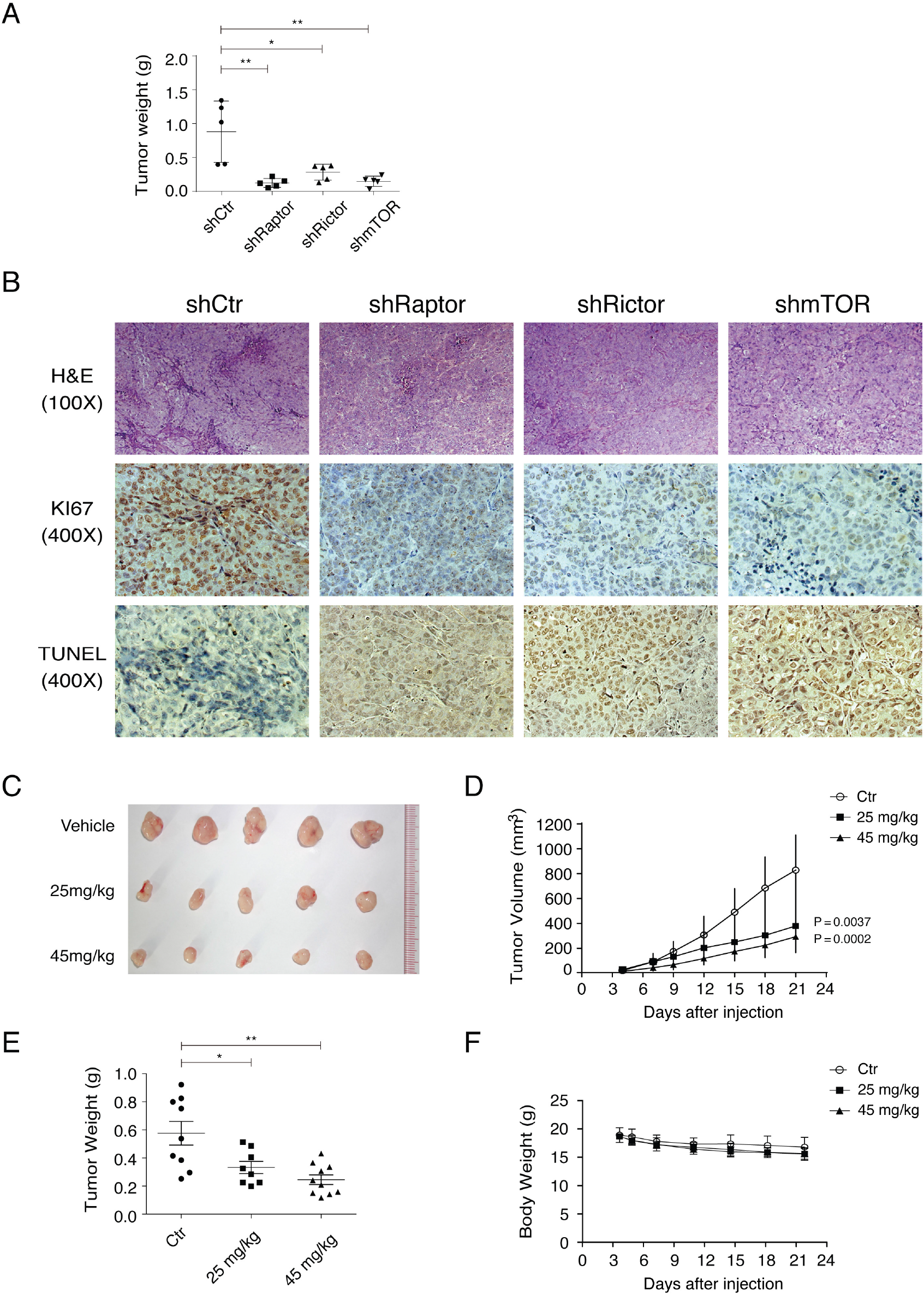
Inhibition of mTOR signaling delayed tumor growth *in vivo*. (**A)** CNE-2 S18 cells (5×10^5^ cells, with Matrigel) and cells in that Raptor, Rictor and mTOR expression had been silenced by specific shRNAs were transplanted subcutaneously into the right flanks of BALB/C-nu/nu mice. After 30 days, tumors were stripped, and tumor mass were weighted. Data are represented as mean ±SD and p-value of one-tailed unpaired t test. (**B)** Tumor sections were examined by H&E staining, IHC for Ki67 and TUNEL staining. (**C)** The effects of mTOR dual inhibitor BEZ235 on NPC tumor formation were examined in NPC xenograft model and mice were treated with BEZ235 in a dose of 45 mg/kg or 25 mg/kg body weight daily by intragastric administration. Images of NPC tumors treated with BEZ235 or vehicles are shown. **(D–F)** The effect of BEZ235 on tumor volume **(D)**, tumor weight **(E)** and body weight of mice **(F)** were illustrated.

In summary, we dissected the roles of distinct mTORC1 and C2 signaling in maintenance of cancer stem cells in NPC. (i) mTORC1 signaling controlled chemotherapy resistance feature of SP population through improving the phosphorylation of 4E-BP1 and increasing ATP-dependent efflux pump ABCG2 in NPC cells. (ii) mTORC2 signaling maintained tumor initiation capability of CSCs through up-regulating c-Myc expression. (iii) Both mTORC1 and mTORC2 can promote the ability of cell migration and invasion of NPC. A model for these functions of the mTOR pathways in regulating nasopharyngeal cancer stem cell characteristics is illustrated in Fig. 7.

**Figure 7.**
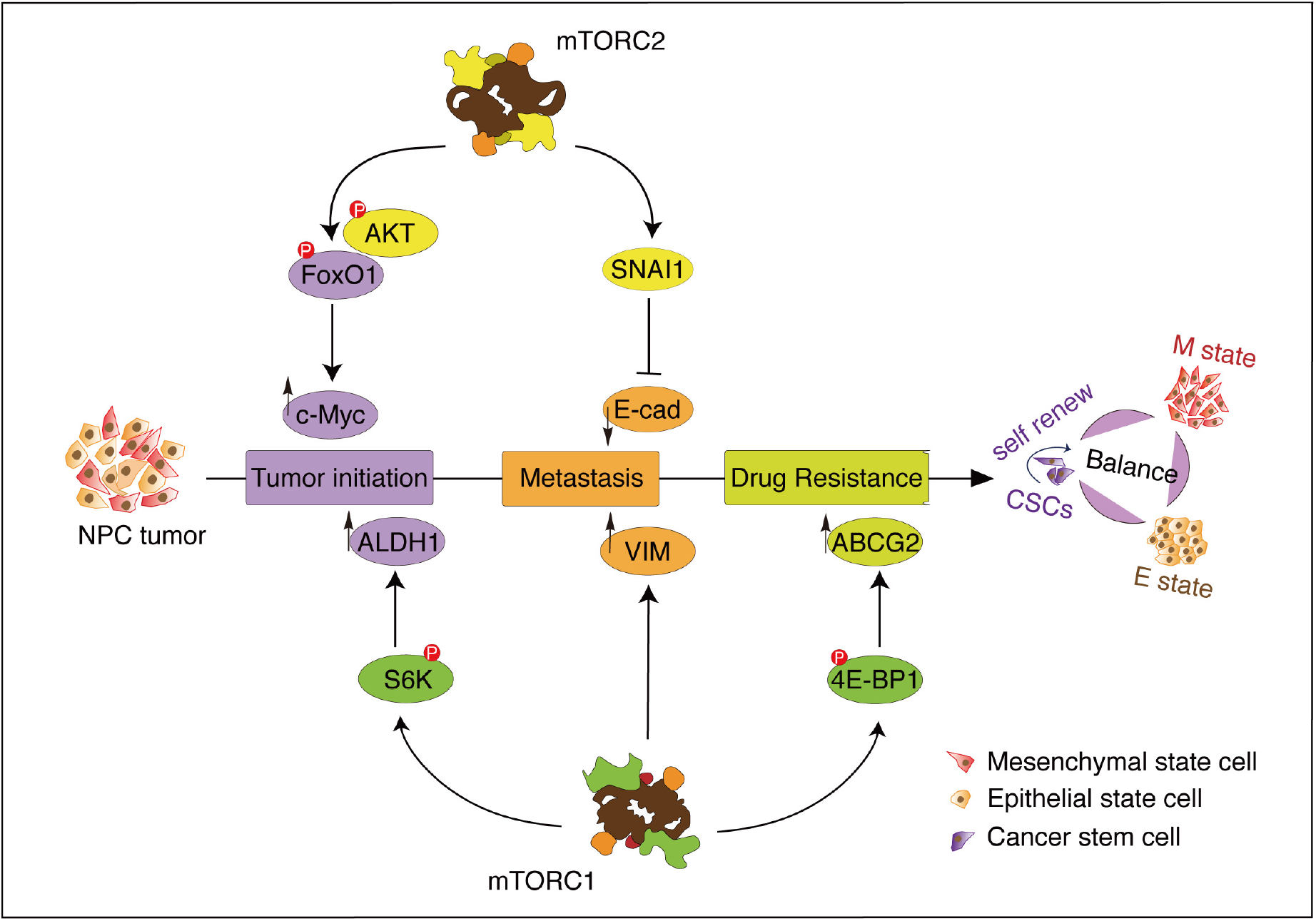
Schematical illustration of a model for mTOR signaling regulating NPC cancer stem cell characteristics.

## Discussion

The vast majority of NPC is the undifferentiated type with invasive and metastatic propensity (40, 41). Recently EMT, which contributes to the generation of cancer stem cells, has emerged as a critical step in cancer initiation, progression, and metastasis (42). Although evidence has suggested that EBV oncogenic proteins LMP1 and LMP2A play crucial roles in the EMT process, as indicated by their up-regulating EMT transcription factor Twist and Snails (5), as well as CSC generation in NPC (43, 44), how LMP1 and LMP2A regulate the EMT process and contribute to each of the CSC tumorigenic properties, such as tumor initiation, migration/invasion, and resistance to anticancer therapies, remains largely unknown. Previous investigations have identified many signal transduction pathways and transcription regulators that can be modulated by LMP1 or LMP2A including TGF-β, mTORC1/NF-κB, ERK-MAPK, and EMT transcription factors twist, snails and Ets1 (9, 45-51). Besides, evidence indicates that mTOR signaling is required for the EMT induction and CSCs maintenance in NPC, and mTORC1 inhibitor rapamycin has a certain degree of inhibitory effect on nasopharyngeal cancer stem cell characteristics (52, 53). These findings compelled us to explore the role of EBV LMPs in mTOR promoting EMT to generate highly tumorigenic NPC cells with cancer stem cell properties. In the current study, we found that LMP1 activated both mTORC1 and mTORC2 signaling pathways that play essential roles in acquiring CSC tumorigenic characteristics, including tumor initiation, metastasis, and therapeutic resistance. Our study showed that mTORC1 and mTORC2 have distinct functions responsible for different tumorigenic properties, respectively.

PI3K-mTOR signaling is one of the most frequently activated pathways in cancer (19), playing roles in tumorigenesis of many aspects including tumor cell proliferation and survival, migration/metastasis and antitumor therapy resistance (20, 21). Recently, the roles of PI3K/mTOR in cancer stem cell establishment have emerged (54-56). Activation of PI3K/mTOR enriched CSCs in breast cancer (55) and blockage of PI3K/mTOR signaling by dual inhibitor VS-5584 preferentially inhibits proliferation and survival of CSCs (56). Our current study confirmed the involvement of mTOR signaling in CSC development in NPC and dissect the roles of distinct signaling (mTORC1 and mTORC2) in the acquisition of different CSC characteristics (schematically illustrated in Fig. 7). (i) Side population (SP), which represents the chemotherapy resistance feature of CSC, was found to be controlled by mTORC1 signaling in NPC. The mTOR signaling pathway has been implicated in multiple anticancer drug resistance mechanisms as many mutations and activation of signaling upstream of mTOR (such as PI3K and AKT) confer drug resistance in various cancers (breast cancer, prostate cancer, etc.) (57). It is likely that the drug resistance property in these cancers is eventually attributed to the mTORC1 function. (ii) Tumor initiation capability is mainly dependent on activated mTORC2 that provides NPC with the capabilities of proliferation and survival through activating c-Myc. c-Myc is an important transcriptional regulator in embryonic stem cells (ESCs), somatic cell reprogramming and cancer. As an unique ESC module, c-Myc drives a transcription program common to ESCs and cancer cells (58). Therefore, mTORC2–c-Myc axis is certainly a primary regulator for maintaining cancer stem cell and tumor initiation property. (iii) Both mTORC1 and mTORC2 can promote the ability of cell migration and invasion of NPC cells. This suggests that mTORC1 and mTORC2 signaling may have overlapped functions or they coordinately control a common pathway. The role of mTORC1 in tumor cell motility, invasion and metastasis has been long recognized in many tumors and cell lines (20, 59-62). Recently, the involvement of mTORC2 in regulation of cell motility and metastasis has also been reported (63, 64). SNAIL, which is known to play an essential role in cell migration, invasion and metastasis and can be positively regulated by mTORC1 via enhancing its translation, was recently found to be controlled by mTORC2 as well through suppressing its ubiquitin-mediated degradation (64). Both mTORC1 and mTORC2 signaling repress E-cadherin via activation of SNAIL. Another important regulatory mechanism in cell migration, invasion and metastasis involves activation of Rho GTPase family (65). Both mTORC1 and mTORC2 regulate the activities of these proteins (63). mTORC1-mediated 4E-BP1 and S6K1 pathways are essential for the expression of some Rho GTPases (RhoA, CDC42 and Rac1), and the actin organization function of Rho GTPases is controlled by mTORC2 (66, 67). The coordinated regulation of NPC tumorigenesis and cancer stem cell generation by mTORC1 and mTORC2 is summarized in the schematic illustration in Fig. 7. Further investigation is warranted to elucidate the detailed mechanisms of how mTOR signaling controls the development of each CSC characteristics in NPC.

The revelation of the roles of mTOR signaling in nasopharyngeal cancer stem cell establishment has implications for therapeutic strategies to treat NPC, especially relapsed and metastatic NPC, by targeting its cancer stem cells using mTOR inhibitors. A thorough comprehension of the roles of different mTOR signaling pathways in distinct tumorigenic features will provide a guideline for designing efficient therapies by choosing specific mTOR inhibitors. For instance, it was shown that mTORC1 inhibitor rapamycin could reduce the side population (SP) cells in breast cancer MCF7 cells (54). However, in our study, rapamycin failed to reduce the SP subpopulation in NPC CEN-2 cells. We believe that this discrepancy results from the activation of the rapamycin-insensitive (RI) mTORC1 pathway in CEN-2 cells; therefore, the inhibitors that can block the RI-mTOR pathway should be used to treat NPC. Indeed, SP cells can be efficiently eliminated when CNE-2 cells were treated with PI3K/mTOR dual inhibitor BEZ-235 (Fig. 4). In addition, given the crucial roles of mTORC2 in regulating tumor initiation, the discovery of new inhibitors, specifically targeting mTORC2, becomes an important and urgent task in new cancer drug development. Recently we identified an mTORC2 potential inhibitor, namely Manassantin B, from the roots of *Saururus chinensis* and found that it can effectively inhibit EBV lytic replication with low cytotoxicity (68). Thus, further study is warranted to evaluate the potential of Manassantin B in inhibition of CSC and the treatment of EBV-associated NPC. Furthermore, our results suggest that cell migration and invasion, which reflects cancer metastatic ability, can be suppressed when both mTORC1 and mTORC2 functions are knocked down or inhibited. Therefore, mTORC1 and mTORC2 dual inhibitors (such as BEZ-235 and VS-5584) could be a better choice for CSC-targeted treatment of NPC to reduce metastasis and achieve durable remission.

## Materials and Methods

### Ethics statement

All animal works were approved by the IACUC of SYSU Zhongshan School of Medicine (No.2017-196). The experiment number is North-D2019-0064. Experiments were carried out under the institutional guidelines of caring laboratory animals, published by the ministry of Science and Technology of People’s Republic of China.

### Cells and chemicals

CNE2-S18 (S18) cells and CNE2-S26 (S26) cells were obtained from Dr. Mu-Sheng Zeng at Sun Yat-sen University Cancer Center, and S18-LMP1 cells and S26-LMP1 cells were established as described previously (6). They were maintained in RPMI 1640 medium supplemented with 5% fetal bovine serum (FBS, Gibco® Life Technologies, #10270-106). Human embryonic kidney (HEK) 293T cells, purchased from American Type Culture Collection (ATCC), were grown in Eagle’s medium (DMEM) supplemented with 10% FBS. All cultures contained 100U/ml penicillin-streptomycin (HyClone Cat# SV30010).

### Antibodies

Antibodies against mTOR signaling pathways have been described in our previous study (68). Antibodies against Vimentin (Cat#5741), Snail (Cat #3879), ZEB1 (Cat #3396), SOX2 (Cat #3579), E-Cadherin (Cat #14472), TWIST1 (Cat #46702), KLF4 (Cat #12173), Nanog (Cat #8822), BMI1(Cat #6964), c-MYC (Cat #18583), Phospho-FoxO1 (Thr24)/FoxO3a (Thr32) (Cat #9464), FoxO1 (Cat #2880), Phospho-4E-BP1 (Thr37/46) (Cat #2855), 4E-BP1 (Cat #9644) were purchased from Cell Signal Technologies. Antibody against EBV Latent Membrane Protein 1 was purchased from abcam (Cat#ab78113).

### Plasmids

Plasmid pcDNA3.1-LMP1 and control vector were kindly provided by Dr. Bijun Huang at Sun Yat-sen University Cancer Center. The pLKO.1-shRNA lentiviral vectors targeting mTOR (Clone ID: NM_004958.2-5477s1c1), Raptor (Clone ID: NM_020761.1-4689s1c1), Rictor (CloneID: NM_152756.2-2620s1c1), as well as pLKO.1-shCTR plasmid, psPAX2 plasmid, and pMD2.G were purchased from Sigma-Aldrich.

### Western blot

Cells were lysed with cell lysis buffer [50 mM Tris-HCl, pH 7.4, 150 mM NaCl, 1% NP-40, 1 mM sodium orthovanadate (Na_3_VO_4_), 20 mM sodium pyrophosphate, 100 mM sodium fluoride, 10% glycerol, protease inhibitor cocktail (1 tablet in 50 mL lysis buffer)]. For nuclear protein detection, RIPA strong lysis buffer (containing 1% NP-40 and 1% Triton X-100) was used. Whole cell lysates were prepared by homogenization and centrifugation at 13,000 rpm for 10 min at 4°C. The whole cell extracts of 50 μg protein was resolved by SDS-PAGE and transferred onto nitrocellulose membranes. The membranes were blocked in 5% non-fat milk in 1×PBS for 1 h, and then incubated in diluted primary antibodies overnight at 4°C. IRDye 680LT and 800CW goat anti-rabbit IgG or anti-mouse IgG antibodies (LI-COR Biosciences) was used as secondary antibody. An Odyssey system (LI-COR) was used for detection of proteins of interest.

### ShRNA-mediated gene silencing

The pLKO.1-shRNA lentiviral plasmids targeting mTOR, Raptor and Rictor, as well as pLKO.1-shCTR lentiviral plasmid, were co-transfected with packaging plasmids psPAX2 and pMD2.G into HEK293T cells. After 72 hours, Media containing lentiviral particles were harvested and subjected to ultracentrifuge to concentrate lentiviruses. S18 cells were transduced with these lentiviruses and selected with 2 μg/ml puromycin for 7 days.

### Immunofluorescence assay (IFA)

Cells were grown on glass coverslips (NEST) for 48 hours. After washing with 1xPBS, cells were fixed using 4% paraformaldehyde for 10 min, permeabilized in 0.1% Triton X-100 for 30 min and blocked in 1%BSA for 1 hour. Then the fixed cells were incubated with anti-Vimentin (1:100 dilution) Antibody for 1 hour at room temperature. Fluor Alexa-555 conjugated anti Rabbit IgG (Life Technologies, 1:200 dilution) was used as secondary antibody. Slides were visualized by Zeiss LSM780 confocal laser scanning system.

### Side population (SP) assay

Cells were digested with 0.25% trypsin, washed twice using 1xPBS, and resuspended in RPMI 1640 medium supplement with 2% FBS in a concentration of 1×10^6^ cells/ml. Hoechst 33342 was added to the cells in a final concentration of 5 μg/ml and incubated for 90 minutes in the dark with periodic mixing at 37°C. Cells were then washed twice with cold 1xPBS and kept on ice for analyzing by BD LSRFortessa.

### Aldehyde dehydrogenase (ALDH) assay

ALDEFLUOR kit (StemCell technologies Cat #01700) was used for identification of cancer stem cells that express high levels of ALDH. Cells were incubated with ALDEFLUOR assay buffer containing ALDEFLUOR reagent (1×10^6^ cells/ml). Cells treated with DEAB reagent was used as a negative control. The samples were incubated at 37°C for 45 min, and then resuspended in ALDEFLUOR assay buffer on ice for analyzing using BD LSRFortessa and CytoFLEX Flow Cytometer.

### Annexin V Apoptosis Assay

S18 cells and cells with Raptor, Rictor or mTOR silenced were collected for apoptosis assay using PE Annexin V Apoptosis Detection Kit I (BD Biosciences Cat#559763). Cells were analyzed using CytoFLEX Flow Cytometer.

### Transwell migration and invasion assays

Cell migration assay and invasion were carried out in 24-well Transwell units (Corning Cat#3422). For a migration assay, 10^5^ cells in 100 μl of serum-free RPMI1640 medium were placed in the top chamber of Transwell. For an invasion assay, 50 μl of diluted matrigel was added to each upper chamber insert of Transwell and the well was incubated at 37°C for 2 hours. Cells (10^5^ in 100 μl of serum-free RPMI1640 medium, starved for 24 hours) were placed in the top chamber of Transwell. The bottom chambers were filled with 600 μl RPMI 1640 medium with 20% FBS. After incubation at 37°C for 20 hours, cells that have passed through the matrigel were fixed with ethanol and stained with crystal violet. The number of migrated cells was counted from multiple randomly selected microscopic visual fields using ImageJ software. Photographs were taken and independent experiments were performed in triplicate.

### MTT assay for cell viability

Cell viability of CNE2 S18 and CNE2 S26 cells after exposed to different chemical compounds was determined by the MTT assay. Cells, seeded into 96-well plates in 200 □l complete RPMI 1640 medium, were treated with Cisplatin, Rapamycin or BEZ235 at various concentrations at 37°C for 72 hours. 20 μl of 5mg/ml MTT [3-(4,5-dimethylthiazol-2-yl)-2,5-diphenyltetrazolium bromide] (Sigma) was added to each well. After 4 hours incubation, the culture medium of each well was discarded, and formazan solubilized in 150 μl DMSO was added. The absorbance of each well was measured at 570 nM. The mean optical density of three wells in treatment groups was used to calculate the percentage of cell viability as follows: Relative cell viability = (A_treatment_ − A_blank_)/(A_control_ − A_blank_) (A = absorbance). The dose response curves were analyzed by GraphPad Prism using the equation “log (inhibitor) vs. response”.

### CFSE cell proliferation assay

S18-shCtr, S18-shRaptor, S18-shRictor and S18-shmTOR cells stained with CellTrace CFSE Cell Proliferation Kit (Invitrogen Cat#C34554) and cultured for 5 days. Cells were analyzed using CytoFLEX Flow Cytometer with 488nm excitation and a 530/30nm emission filter.

### Tumorsphere formation assay

Tumorsphere formation assay were carried out in ultra-low attachment multiple well plate (Corning® Costar® Cat#CLS3471). 100 cells in 2 ml of tumorsphere medium (serum-free DMEM/F12 medium supplemented with B27, 20 μg/ml EGF and 10 μg/ml bFGF) were loaded into each well. After 10 days incubation, tumorspheres were counted and visualized by Zeiss cell observe Z1. Experiments were carried out in triplicate with at least three replicates per concentration.

### Cell migration assay

Cell migration was measured using the wound healing assay. Cells were seeded into 12-well plates in a density of 2×10^5^ cells per well and cultured until confluent. The cell monolayer was scratched using a yellow pipette tip and washed twice with 1x PBS then changed the medium to serum free RPMI 1640 medium. Initial images of three independent areas after each scratch were acquired at time zero. Images of the same area were captured again after incubation at 37°C for 20 hours.

### Tumor xenograft experiment

BALB/C-nu/nu mice were purchased from the Laboratory Animal Center of Sun Yat-Sen University. Tumor cells were suspended in 200 μl of diluted matrigel (1:8 diluted with PBS) and inoculated subcutaneously into the right flanks of 4-5 weeks old mice at 5×10^5^ cells/animal. 3 days after injection, mice were randomly deviled into three groups and treated with vehicles or BEZ235 in a dose of 25 or 45 mg/kg by daily intragastric administration. The tumor volumes were measured using a vernier caliper every other day. The tumor weights were measured at 20 days after injection.

### Histological analyses

Tissue processing, Hematoxylin and eosin (H&E) staining and Immunohistochemistry (IHC) staining were performed as described previously (69). TUNEL staining was performed using the DeadEnd™ Colorimetric TUNEL System (Promega, Part Numbers: G7130 and G7360) per the manufacture’s protocol. Photomicrographs were acquired using a Zeiss cell observe Z1.

### Statistical Analysis

The Student’s unpaired t-test was used to compare the data from two study groups. A P value <0.05 (* denotes a P value<0.05, show 4 significant digits) was used to determine statistical significance. Error bars represent the SD (Standard Deviation). Data was analyzed by GraphPad Prism.

## Acknowledgment

We thank all members of Yuan Lab for critical reading of this manuscript and helpful discussion. This study was supported by the National Natural Science Foundation of China (NSFC 81772177).

## References

1. Young LS, Dawson CW. 2014. Epstein-Barr virus and nasopharyngeal carcinoma. Chin J Cancer 33:581–90.

2. Pathmanathan R, Prasad U, Chandrika G, Sadler R, Flynn K, Raab-Traub N. 1995. Undifferentiated, nonkeratinizing, and squamous cell carcinoma of the nasopharynx. Variants of Epstein-Barr virus-infected neoplasia. Am J Pathol 146:1355–67.

3. Sun X, Su S, Chen C, Han F, Zhao C, Xiao W, et al. 2014. Long-term outcomes of intensity-modulated radiotherapy for 868 patients with nasopharyngeal carcinoma: an analysis of survival and treatment toxicities. Radiother Oncol 110:398–403.

4. Tsao SW, Tramoutanis G, Dawson CW, Lo AK, Huang DP. 2002. The significance of LMP1 expression in nasopharyngeal carcinoma. Semin Cancer Biol 12:473–87.

5. Horikawa T, Yang J, Kondo S, Yoshizaki T, Joab I, Furukawa M, et al. 2007. Twist and epithelial-mesenchymal transition are induced by the EBV oncoprotein latent membrane protein 1 and are associated with metastatic nasopharyngeal carcinoma. Cancer Res 67:1970–8.

6. Zhu N, Xu X, Wang Y, Zeng MS, Yuan Y. 2021. EBV latent membrane proteins promote hybrid epithelial-mesenchymal and extreme mesenchymal states of nasopharyngeal carcinoma cells for tumorigenicity. PLoS Pathog 17:e1009873.

7. Xiang T, Lin YX, Ma W, Zhang HJ, Chen KM, He GP, et al. 2018. Vasculogenic mimicry formation in EBV-associated epithelial malignancies. Nat Commun 9:5009.

8. Chen J, Hu CF, Hou JH, Shao Q, Yan LX, Zhu XF, et al. 2010. Epstein-Barr virus encoded latent membrane protein 1 regulates mTOR signaling pathway genes which predict poor prognosis of nasopharyngeal carcinoma. J Transl Med 8:30.

9. Zhang J, Jia L, Lin W, Yip YL, Lo KW, Lau VMY, et al. 2017. Epstein-Barr Virus-Encoded Latent Membrane Protein 1 Upregulates Glucose Transporter 1 Transcription via the mTORC1/NF-κB Signaling Pathways. J Virol 91:e02168–16.

10. Saxton RA, Sabatini DM. 2017. mTOR Signaling in Growth, Metabolism, and Disease. Cell 168:960–976.

11. Sabatini DM. 2006. mTOR and cancer: insights into a complex relationship. Nat Rev Cancer 6:729–34.

12. Kim DH, Sarbassov DD, Ali SM, King JE, Latek RR, Erdjument-Bromage H, et al. 2002. mTOR interacts with raptor to form a nutrient-sensitive complex that signals to the cell growth machinery. Cell 110:163–75.

13. Sarbassov DD, Ali SM, Kim DH, Guertin DA, Latek RR, Erdjument-Bromage H, et al. 2004. Rictor, a novel binding partner of mTOR, defines a rapamycin-insensitive and raptor-independent pathway that regulates the cytoskeleton. Curr Biol 14:1296–302.

14. Frias MA, Thoreen CC, Jaffe JD, Schroder W, Sculley T, Carr SA, et al. 2006. mSin1 is necessary for Akt/PKB phosphorylation, and its isoforms define three distinct mTORC2s. Curr Biol 16:1865–70.

15. Zinzalla V, Stracka D, Oppliger W, Hall MN. 2011. Activation of mTORC2 by association with the ribosome. Cell 144:757–68.

16. Gaubitz C, Prouteau M, Kusmider B, Loewith R. 2016. TORC2 Structure and Function. Trends Biochem Sci 41:532–545.

17. Su B, Jacinto E. 2011. Mammalian TOR signaling to the AGC kinases. Crit Rev Biochem Mol Biol 46:527–47.

18. Sarbassov DD, Guertin DA, Ali SM, Sabatini DM. 2005. Phosphorylation and Regulation of Akt/PKB by the Rictor-mTOR Complex. Science 307:1098–1101.

19. Zhang Y, Kwok-Shing Ng P, Kucherlapati M, Chen F, Liu Y, Tsang YH, et al. 2017. A Pan-Cancer Proteogenomic Atlas of PI3K/AKT/mTOR Pathway Alterations. Cancer Cell 31:820–832.e3.

20. Hsieh AC, Liu Y, Edlind MP, Ingolia NT, Janes MR, Sher A, et al. 2012. The translational landscape of mTOR signalling steers cancer initiation and metastasis. Nature 485:55–61.

21. Weiler M, Blaes J, Pusch S, Sahm F, Czabanka M, Luger S, et al. 2014. mTOR target NDRG1 confers MGMT-dependent resistance to alkylating chemotherapy. Proc Natl Acad Sci U S A 111:409–14.

22. Qian CN, Berghuis B, Tsarfaty G, Bruch M, Kort EJ, Ditlev J, et al. 2006. Preparing the “soil”: the primary tumor induces vasculature reorganization in the sentinel lymph node before the arrival of metastatic cancer cells. Cancer Res 66:10365–76.

23. Ginestier C, Hur MH, Charafe-Jauffret E, Monville F, Dutcher J, Brown M, et al. 2007. ALDH1 is a marker of normal and malignant human mammary stem cells and a predictor of poor clinical outcome. Cell Stem Cell 1:555–67.

24. Li W, Ma H, Zhang J, Zhu L, Wang C, Yang Y. 2017. Unraveling the roles of CD44/CD24 and ALDH1 as cancer stem cell markers in tumorigenesis and metastasis. Sci Rep 7:13856.

25. Richard V, Nair MG, Santhosh Kumar TR, Pillai MR. 2013. Side population cells as prototype of chemoresistant, tumor-initiating cells. Biomed Res Int 2013:517237–517237.

26. Wang J, Guo LP, Chen LZ, Zeng YX, Lu SH. 2007. Identification of cancer stem cell-like side population cells in human nasopharyngeal carcinoma cell line. Cancer Res 67:3716–24.

27. Feldman ME, Apsel B, Uotila A, Loewith R, Knight ZA, Ruggero D, Shokat KM. 2009. Active-site inhibitors of mTOR target rapamycin-resistant outputs of mTORC1 and mTORC2. PLoS Biol 7:e38.

28. Zhou S, Schuetz JD, Bunting KD, Colapietro AM, Sampath J, Morris JJ, et al. 2001. The ABC transporter Bcrp1/ABCG2 is expressed in a wide variety of stem cells and is a molecular determinant of the side-population phenotype. Nat Med 7:1028–34.

29. Dou J, Jiang C, Wang J, Zhang X, Zhao F, Hu W, et al. 2011. Using ABCG2-molecule-expressing side population cells to identify cancer stem-like cells in a human ovarian cell line. Cell Biol Int 35:227–34.

30. Taylor NMI, Manolaridis I, Jackson SM, Kowal J, Stahlberg H, Locher KP. 2017. Structure of the human multidrug transporter ABCG2. Nature 546:504–509.

31. Bleau AM, Hambardzumyan D, Ozawa T, Fomchenko EI, Huse JT, Brennan CW, Holland EC. 2009. PTEN/PI3K/Akt pathway regulates the side population phenotype and ABCG2 activity in glioma tumor stem-like cells. Cell Stem Cell 4:226–35.

32. Ponti D, Costa A, Zaffaroni N, Pratesi G, Petrangolini G, Coradini D, et al. 2005. Isolation and in vitro propagation of tumorigenic breast cancer cells with stem/progenitor cell properties. Cancer Res 65:5506–11.

33. Koh SP, Brasch HD, de Jongh J, Itinteang T, Tan ST. 2019. Cancer stem cell subpopulations in moderately differentiated head and neck cutaneous squamous cell carcinoma. Heliyon 5:e02257.

34. Müller M, Hermann PC, Liebau S, Weidgang C, Seufferlein T, Kleger A, Perkhofer L. 2016. The role of pluripotency factors to drive stemness in gastrointestinal cancer. Stem Cell Res 16:349–57.

35. Wilson A, Murphy MJ, Oskarsson T, Kaloulis K, Bettess MD, Oser GM, et al. 2004. c-Myc controls the balance between hematopoietic stem cell self-renewal and differentiation. Genes Dev 18:2747–63.

36. Peck B, Ferber EC, Schulze A. 2013. Antagonism between FOXO and MYC Regulates Cellular Powerhouse. Front Oncol 3:96.

37. Brunet A, Bonni A, Zigmond MJ, Lin MZ, Juo P, Hu LS, et al. 1999. Akt promotes cell survival by phosphorylating and inhibiting a Forkhead transcription factor. Cell 96:857–68.

38. Wilhelm K, Happel K, Eelen G, Schoors S, Oellerich MF, Lim R, et al. 2016. FOXO1 couples metabolic activity and growth state in the vascular endothelium. Nature 529:216–20.

39. Gabay M, Li Y, Felsher DW. 2014. MYC activation is a hallmark of cancer initiation and maintenance. Cold Spring Harb Perspect Med 4:a014241.

40. Lo KW, To KF, Huang DP. 2004. Focus on nasopharyngeal carcinoma. Cancer Cell 5:423–8.

41. Tao Q, Chan AT. 2007. Nasopharyngeal carcinoma: molecular pathogenesis and therapeutic developments. Expert Rev Mol Med 9:1–24.

42. Shibue T, Weinberg RA. 2017. EMT, CSCs, and drug resistance: the mechanistic link and clinical implications. Nat Rev Clin Oncol 14:611–629.

43. Kondo S, Wakisaka N, Muramatsu M, Zen Y, Endo K, Murono S, et al. 2011. Epstein-Barr virus latent membrane protein 1 induces cancer stem/progenitor-like cells in nasopharyngeal epithelial cell lines. J Virol 85:11255–64.

44. Kong QL, Hu LJ, Cao JY, Huang YJ, Xu LH, Liang Y, et al. 2010. Epstein-Barr virus-encoded LMP2A induces an epithelial-mesenchymal transition and increases the number of side population stem-like cancer cells in nasopharyngeal carcinoma. PLoS Pathog 6:e1000940.

45. Kim KR, Yoshizaki T, Miyamori H, Hasegawa K, Horikawa T, Furukawa M, et al. 2000. Transformation of Madin-Darby canine kidney (MDCK) epithelial cells by Epstein-Barr virus latent membrane protein 1 (LMP1) induces expression of Ets1 and invasive growth. Oncogene 19:1764–71.

46. Kang Y, Massagué J. 2004. Epithelial-mesenchymal transitions: twist in development and metastasis. Cell 118:277–9.

47. Dawson CW, Laverick L, Morris MA, Tramoutanis G, Young LS. 2008. Epstein-Barr virus-encoded LMP1 regulates epithelial cell motility and invasion via the ERK-MAPK pathway. J Virol 82:3654–64.

48. Morris MA, Young LS, Dawson CW. 2008. DNA tumour viruses promote tumour cell invasion and metastasis by deregulating the normal processes of cell adhesion and motility. Eur J Cell Biol 87:677–97.

49. Sides MD, Klingsberg RC, Shan B, Gordon KA, Nguyen HT, Lin Z, et al. 2011. The Epstein-Barr virus latent membrane protein 1 and transforming growth factor--β1 synergistically induce epithelial--mesenchymal transition in lung epithelial cells. Am J Respir Cell Mol Biol 44:852–62.

50. Horikawa T, Yang J, Kondo S, Yoshizaki T, Joab I, Furukawa M, et al. 2007. Twist and Epithelial-Mesenchymal Transition Are Induced by the EBV Oncoprotein Latent Membrane Protein 1 and Are Associated with Metastatic Nasopharyngeal Carcinoma. Cancer Res 67:1970–8.

51. Horikawa T, Yoshizaki T, Kondo S, Furukawa M, Kaizaki Y, Pagano JS. 2011. Epstein-Barr Virus latent membrane protein 1 induces Snail and epithelial–mesenchymal transition in metastatic nasopharyngeal carcinoma. Br J Cancer 104:1160–7.

52. Wang MH, Sun R, Zhou XM, Zhang MY, Lu JB, Yang Y, et al. 2018. Epithelial cell adhesion molecule overexpression regulates epithelial-mesenchymal transition, stemness and metastasis of nasopharyngeal carcinoma cells via the PTEN/AKT/mTOR pathway. Cell Death Dis 9:2.

53. Yang C, Zhang Y, Zhang Y, Zhang Z, Peng J, Li Z, et al. 2015. Downregulation of cancer stem cell properties via mTOR signaling pathway inhibition by rapamycin in nasopharyngeal carcinoma. Int J Oncol 47:909–17.

54. Zhou J, Wulfkuhle J, Zhang H, Gu P, Yang Y, Deng J, et al. 2007. Activation of the PTEN/mTOR/STAT3 pathway in breast cancer stem-like cells is required for viability and maintenance. Proc Natl Acad Sci U S A 104:16158–63.

55. Korkaya H, Paulson A, Charafe-Jauffret E, Ginestier C, Brown M, Dutcher J, et al. 2009. Regulation of mammary stem/progenitor cells by PTEN/Akt/beta-catenin signaling. PLoS Biol 7:e1000121.

56. Kolev VN, Wright QG, Vidal CM, Ring JE, Shapiro IM, Ricono J, et al. 2015. PI3K/mTOR dual inhibitor VS-5584 preferentially targets cancer stem cells. Cancer Res 75:446–55.

57. Jiang BH, Liu LZ. 2008. Role of mTOR in anticancer drug resistance: perspectives for improved drug treatment. Drug Resist Updat 11:63–76.

58. Kim J, Woo AJ, Chu J, Snow JW, Fujiwara Y, Kim CG, et al. 2010. A Myc network accounts for similarities between embryonic stem and cancer cell transcription programs. Cell 143:313–24.

59. Berven LA, Willard FS, Crouch MF. 2004. Role of the p70(S6K) pathway in regulating the actin cytoskeleton and cell migration. Exp Cell Res 296:183–95.

60. Liu L, Li F, Cardelli JA, Martin KA, Blenis J, Huang S. 2006. Rapamycin inhibits cell motility by suppression of mTOR-mediated S6K1 and 4E-BP1 pathways. Oncogene 25:7029–40.

61. Lamouille S, Derynck R. 2007. Cell size and invasion in TGF-beta-induced epithelial to mesenchymal transition is regulated by activation of the mTOR pathway. J Cell Biol 178:437–51.

62. Gomez-Cambronero J. 2003. Rapamycin inhibits GM-CSF-induced neutrophil migration. FEBS Lett 550:94–100.

63. Gulhati P, Bowen KA, Liu J, Stevens PD, Rychahou PG, Chen M, et al. 2011. mTORC1 and mTORC2 regulate EMT, motility, and metastasis of colorectal cancer via RhoA and Rac1 signaling pathways. Cancer Res 71:3246–56.

64. Zhang S, Qian G, Zhang QQ, Yao Y, Wang D, Chen ZG, et al. 2019. mTORC2 Suppresses GSK3-Dependent Snail Degradation to Positively Regulate Cancer Cell Invasion and Metastasis. Cancer Res 79:3725–3736.

65. Hall A. 1998. Rho GTPases and the actin cytoskeleton. Science 279:509–14.

66. Liu L, Luo Y, Chen L, Shen T, Xu B, Chen W, et al. 2010. Rapamycin inhibits cytoskeleton reorganization and cell motility by suppressing RhoA expression and activity. J Biol Chem 285:38362–73.

67. Jacinto E, Loewith R, Schmidt A, Lin S, Rüegg MA, Hall A, et al. 2004. Mammalian TOR complex 2 controls the actin cytoskeleton and is rapamycin insensitive. Nat Cell Biol 6:1122–8.

68. Wang Q, Zhu N, Hu J, Wang Y, Xu J, Gu Q, et al. 2020. The mTOR inhibitor manassantin B reveals a crucial role of mTORC2 signaling in Epstein-Barr virus reactivation. J Biol Chem 295:7431–7441.

69. Li Y, Zhong C, Liu D, Yu W, Chen W, Wang Y, et al. 2018. Evidence for Kaposi Sarcoma Originating from Mesenchymal Stem Cell through KSHV-induced Mesenchymal-to-Endothelial Transition. Cancer Res 78:230–245.

